# Unified and standardized mass spectrometry data processing in Python using spectrum_utils

**DOI:** 10.1101/2022.10.04.510894

**Authors:** Wout Bittremieux, Lev Levitsky, Matteo Pilz, Timo Sachsenberg, Florian Huber, Mingxun Wang, Pieter C. Dorrestein

## Abstract

spectrum_utils is a Python package for mass spectrometry data processing and visualization. Since its introduction, spectrum_utils has grown into a fundamental software solution that powers various applications in proteomics and metabolomics, ranging from spectrum preprocessing prior to spectrum identification and machine learning applications, to spectrum plotting from online data repositories, and assisting data analysis tasks for dozens of other projects. Here we present updates to spectrum_utils, which include new functionality to integrate mass spectrometry community data standards, enhanced mass spectral data processing, and unified mass spectral data visualization in Python. spectrum_utils is freely available as open source at https://github.com/bittremieux/spectrum_utils.

## Introduction

There exists a rich ecosystem of open-source mass spectrometry (MS) software. Some of the most popular MS software libraries in common programming languages, such as Python and R, include Pyteomics,^1,2^ pyOpenMS,^3^ matchms,^4^ pymzml,^5,6^ and MSnbase.^7,8^ Compared to vendor software and other closed-source MS software, these open-source solutions provide flexibility to develop powerful functionalities, are verifiable through their open-source nature, and have garnered widespread community support to grow their capabilities. Although each library predominantly covers its unique use cases, there are common tasks that frequently occur, such as MS/MS spectrum processing and visualization.

In its first release, spectrum_utils^9^ provided a lightweight solution to cover common MS data manipulation functionalities. Its high-level application programming interface (API) allows developers to quickly prototype computational ideas for mass spectrometry projects. The data processing functionality is optimized for computational efficiency to handle large volumes of data (hundreds to thousands of files) that are generated during MS experiments. This dovetails with visualization functionality to produce publication-quality and interactive spectrum graphics. Since its introduction, spectrum_utils has been used in tools that perform spectral library searching^10^ and spectrum clustering,^11^ to preprocess MS/MS spectra prior to deep learning applications,^12,13^ to plot MS/MS spectra from online data repositories,^14^ and to assist in MS/MS processing and visualization efforts for dozens of other projects.^15–22^

Here we present recent updates to spectrum_utils. It has now been extended with functionality for common and versatile data manipulation and visualization tasks when working with MS data in Python, including support for community data standards, updated visualization functionalities, and performance improvements. This has enabled spectrum_utils to grow into a building block of the MS Python ecosystem. It functions both as a client, to provide basic functionality for other software tools and by accessing external web resources, and as a server, through integration in key online tools for end users. With a strong focus on open source, compatibility and integration with other software libraries, and support of community standards, spectrum_utils is contributing to make MS data and software open, accessible, and reusable for the entire community. spectrum_utils is freely available as open source under the Apache 2.0 license at https://github.com/bittremieux/spectrum_utils.

## Methods & Results

### Supporting mass spectrometry community data standards for data access, processing, and visualization

spectrum_utils supports several official data standards and best practices developed by the Proteomics Standards Initiative (PSI) of the Human Proteome Organization (HUPO).^23^ Specifically, spectrum_utils has now integrated support for the Universal Spectrum Identifier (USI)^24^ for convenient retrieval of MS data from ProteomeXchange^25^ and other online resources, the ProForma 2.0 specification^26^ to encode proteoform information, and standardized fragment ion annotations (https://github.com/HUPO-PSI/mzSpecLib/).

The Universal Spectrum Identifier^24^ is a mechanism that provides unique identifiers for MS data present in ProteomeXchange repositories. Rather than only including spectral evidence of novel discoveries as static figures in scientific manuscripts or being relegated to supporting information, USIs make MS data Findable, Accessible, Interoperable, and Reusable (FAIR). spectrum_utils supports the USI specification to retrieve proteomics mass spectra from ProteomeXchange resources, including PRIDE,^27^ MassIVE, PeptideAtlas,^28^ and jPOST,^29^ through integration with Pyteomics,^1,2^ as well as metabolomics mass spectra from GNPS/MassIVE,^30^ MassBank,^31^ MetaboLights,^32^ Metabolomics Workbench,^33^ and MS2LDA,^34^ through integration with the GNPS’ Metabolomics USI Resolver.^14^ USI integration in spectrum_utils makes it possible to effortlessly load MS data from virtually all major proteomics and metabolomics data platforms and start processing it in Python using a single line of code (**Figure 1a**). Furthermore, as the Metabolomics USI resolver provides a convenient web API to programmatically retrieve MS data, indirectly spectrum_utils also facilitates the loading of MS data in any programming language.

**Figure 1:**
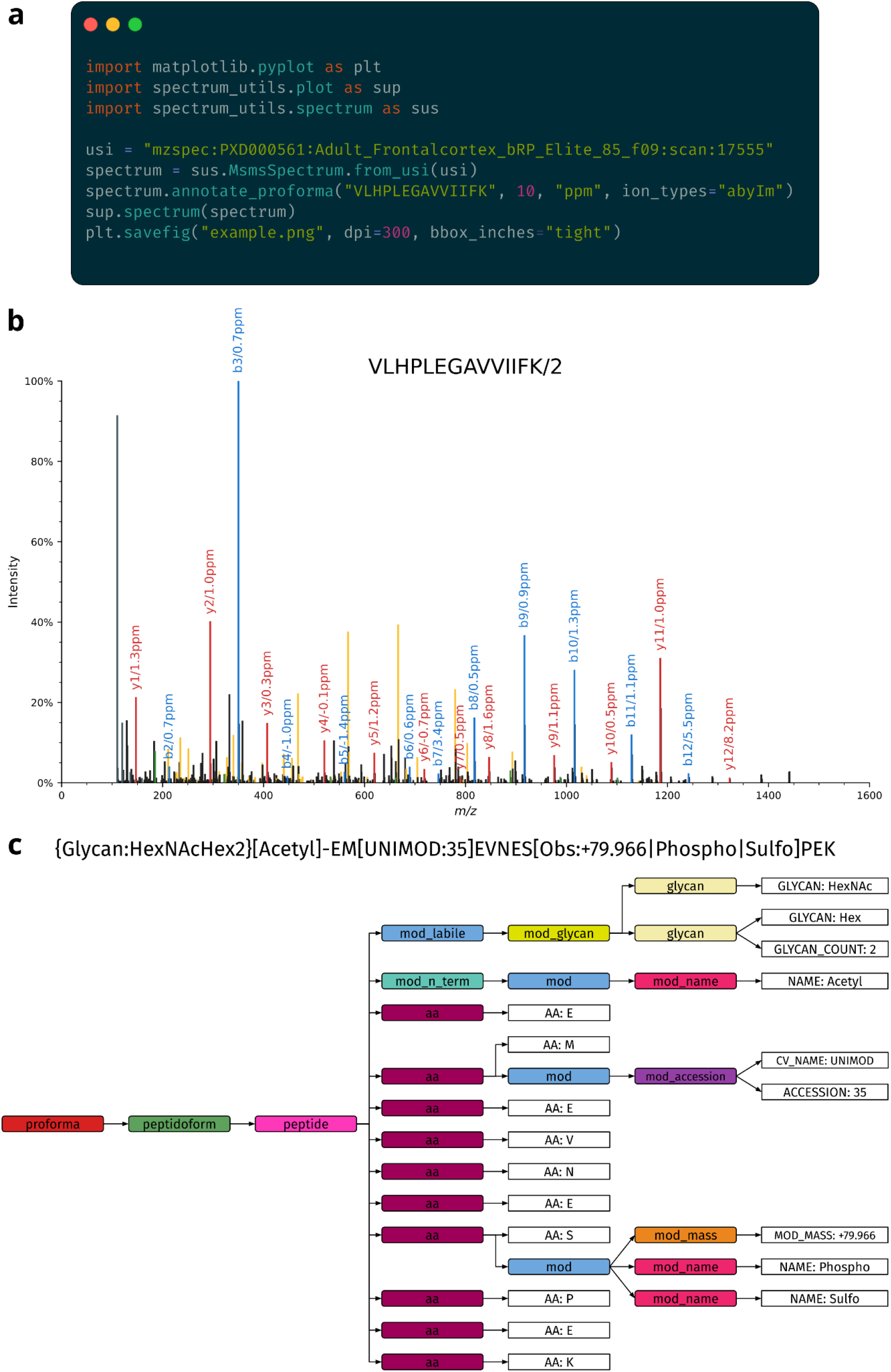
**(a)** spectrum_utils provides a convenient and high-level interface to process MS data. Fully functional code example for loading an MS/MS spectrum using its uSi, annotating the spectrum with the corresponding peptide sequence using the ProForma 2.0 format, and visualizing the spectrum using only a small number of lines of code. **(b)** Spectrum plot corresponding to the code example in (a). The final code was adapted only minimally for visualization purposes. spectrum_utils can annotate known peaks, including typical b and y peptide fragments (highlighted), as well as internal fragment ions (yellow), and immonium ions (dark gray). Different information for annotated peaks, such as the ion type and deviation from the theoretical fragment *m/z*, can be reported. **(c)** Internal representation of the abstract syntax tree when parsing a complex ProForma string that contains several modifications, including labile glycan modifications, modifications specified by their CV accession or CV name, and an observed modification mass of +79.966 that can be interpreted as a phosphorylation or a sulfation.

The ProForma 2.0 format^26^ is a compact way to encode peptidoforms and proteoforms by unambiguously specifying post-translational modifications (PTMs), which was recently developed for use in both bottom-up and top-down proteomics through joint efforts of the PSI and the Consortium for Top-Down Proteomics. ProForma 2.0 specifies several levels of compliance, including base-level support, top-down extensions, cross-linking extensions, glycan extensions, and more. spectrum_utils uniquely provides full support for all ProForma 2.0 levels, including all optional extensions, in a convenient Python interface (**Figure 1a**). Internally, spectrum_utils represents the ProForma 2.0 specification as a formal grammar which is used to create an abstract syntax tree when parsing a ProForma string (**Figure 1c**). This approach is similar to how compilers interpret complex source code instructions, and the formal grammar is the only existing codified representation for ProForma 2.0 that is machine-readable. This is an extremely robust and scalable solution to cover the full ProForma 2.0 specification, including optional extensions and edge cases, compared to alternative approaches, such as combinations of regular expressions. To interpret nodes in the abstract syntax tree that represent PTMs, spectrum_utils supports all controlled vocabularies (CVs) relevant for ProForma 2.0, including Unimod,^35^ PSI-MOD,^36^ RESID, XL-MOD,^37^ and the Glycan Naming Ontology,^38^ as well as PTMs specified by observed precursor mass differences. While traversing an abstract syntax tree, upon the first occurrence of a CV term, spectrum_utils will automatically download the corresponding CV file from its online resource and parse the CV term by its name or accession. Additionally, spectrum_utils will store the CV files in a local cache to minimize the time required for further lookups. Retrieving CV terms from a remote or local CV file is performed in a way that is completely transparent to the user.

When annotating an MS/MS spectrum with its ProForma string, spectrum_utils calculates all theoretical fragments and annotates matching fragment ions. spectrum_utils can consider a wide variety of theoretical fragments, including the primary a, b, c, x, y, and z peptide fragments; internal fragment ions, which result from two amide bond cleavages and thus do not contain either terminus; immonium ions, which are internal fragments for individual amino acids formed by a b/y cleavage on the N-terminal side and an a/x cleavage on the C-terminal side; intact precursor ions; reporter ions from isobaric labeling; and any of the preceding ions that have undergone a neutral loss, such as loss of hydrogen, ammonia, water, etc. Next, spectrum_utils can highlight all of the fragment ions with their annotations while plotting the MS/MS spectrum to visually validate the peptide-spectrum match (**Figure 1b**).

### Case study: Annotating peaks in tandem mass spectra

We have used spectrum_utils to annotate fragment ions for 2,153,703 MS/MS spectra in the MassIVE-KB v1 spectral library.^39^ This case study exemplifies how the powerful functionality to annotate spectra with ProForma peptide information and peak interpretation by matching observed fragment ions can be used. MassIVE-KB is a repository-wide human higher-energy collisional dissociation (HCD) spectral library, derived from over 30 TB of human MS/MS data from 227 public proteomics datasets on the MassIVE repository. Peptide identifications for MassIVE-KB were previously obtained using MSGF+^40^ to search the spectra against the UniProt^41^ human reference proteome database (v. 23 May 2016). Cysteine carbamidomethylation was set as a fixed modification, and variable modifications were methionine oxidation, N-terminal acetylation, N-terminal carbamylation, pyroglutamate formation from glutamine, and deamidation of asparagine and glutamine. MSGF+ was configured to allow one ^13^C precursor mass isotope, at most one non-tryptic terminus, and 10 ppm precursor mass tolerance. The searches were individually filtered at 1% PSM-level FDR.

To interpret the observed fragment ions, we considered a, b, c, x, y, and z peptide fragments, immonium ions, internal fragment ions, and intact precursor ions. Additionally, the following neutral losses were considered for any of these ions: loss of hydrogen (H), loss of ammonia (NH3), loss of water (H2O), loss of carbon monoxide (CO), loss of carbon dioxide (CO2), loss of formamide (HCONH2), loss of formic acid (HCOOH), loss of methanesulfenic acid (CH4OS), loss of sulfur trioxide (SO3), loss of metaphosphoric acid (HPO3), loss of mercaptoacetamide (C2H5NOS), loss of mercaptoacetic acid (C2H4O2S), and loss of phosphoric acid (H3PO4). Fragments with charges up to the spectrum precursor charge were considered, and fragment ions were assigned a peak interpretation if their observed *m/z* was within 20 ppm of the theoretical *m*/*z*. In case multiple peak interpretations matched an observed fragment ion, heuristics were used to select the most relevant peak annotation by preferring y and b fragments, as well as fragments without a neutral loss.

Despite the many theoretical fragments that are possible, peak annotation of over 2 million MS/MS spectra took under 2.5 hours using an Intel Xeon Gold 6148 processor (2.4 GHz, 40 cores). On average, 74% of the observed intensity of the spectra could be explained by a matching peak interpretation. As expected for HCD data, the most prevalent ion types were y ions, explaining 33% of the fragment intensities on average, while b ions explained 13% of the fragment intensities (**Figure 2a**). Internal fragment ions also covered a non-negligible amount of intensity. Although the average intensity of individual internal fragment ions was third-lowest (after x and z ions), their cumulative explained intensity amounted to 15% on average due to the very large number of small peaks that could be interpreted as internal fragments.

**Figure 2:**
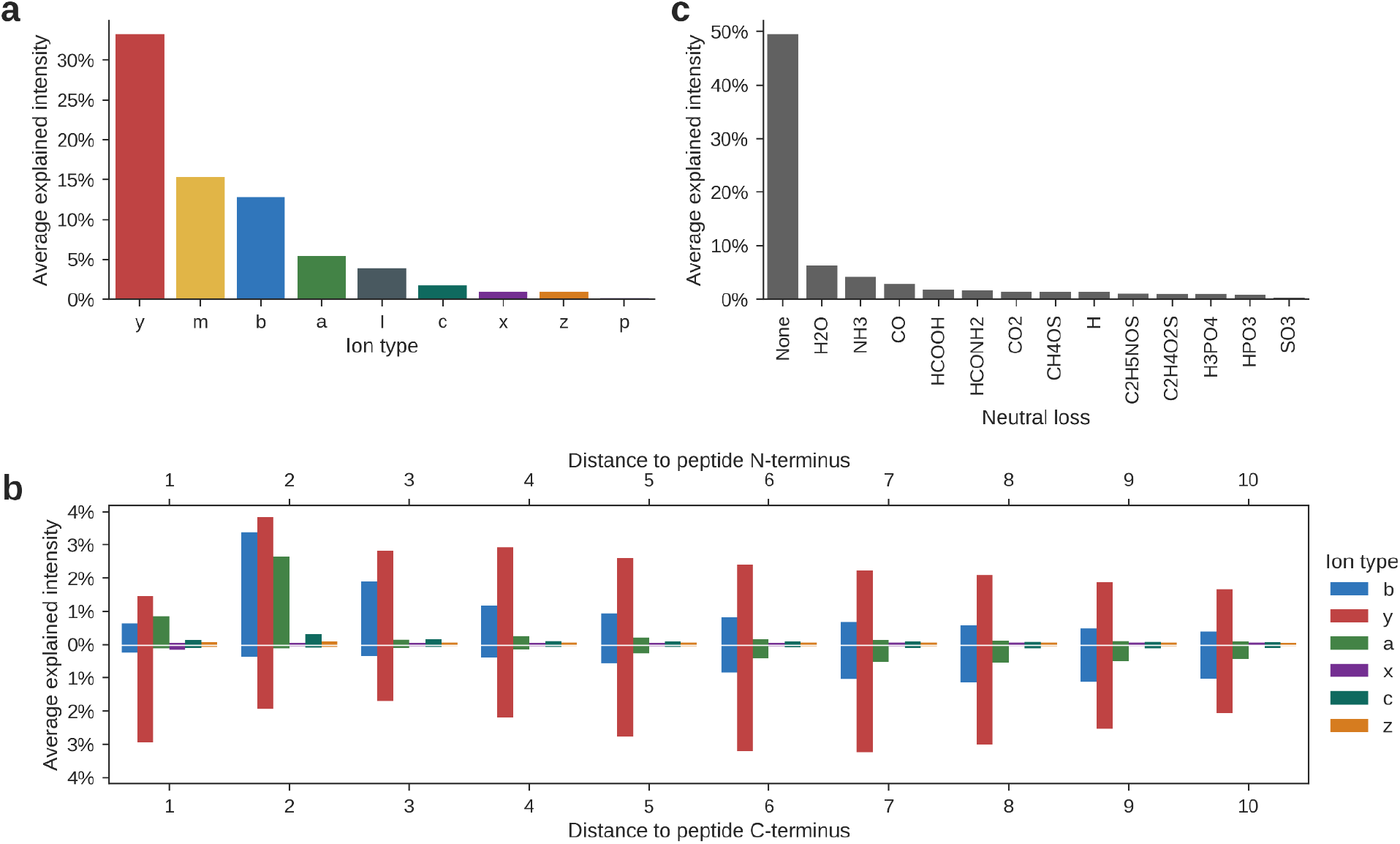
**(a)** Average explained intensity for fragment ions across 2 million HCD MS/MS spectra from the MassIVE-KB spectral library. Considered ion types are peptide fragments “a”, “b”, “c”, “x”, “y”, and “z”; internal fragment ions “m”, immonium ions “I”, and intact precursor ions “p”. **(b)** Intensity of peptide fragments related to their location in the peptides. For peptides with length *n*, distances to the peptide N-terminus correspond to b_1_, b_2_, … and y_*n*-1_, y_*n*-2_, … fragments (top), and distances to the peptide C-terminus correspond to b_*n*-1_, b_*n*___2_, … and y_1_, y_2_, … fragments (bottom). Distances for a/c and x/z fragments are interpreted equivalently. **(c)** Average explained intensity for neutral losses of the interpreted fragment ions.

When further examining the intensity of peptide fragments based on their location in the peptides (**Figure 2b**), we can observe that y ions provide the dominant signal across a wide range of fragment locations, followed by b ions, which are most intense in locations near the peptide N-terminus. Additionally, peptide a ions commonly occur for small N-terminal fragments. Thus, even though sequence database search engines often only consider b and y ions for HCD data, taking small a ions into account might help bolster confidence in the spectrum annotations.

Although the majority of explained intensity corresponds to fragments that do not include a neutral loss, a quarter of the observed intensity matches fragment ions that have undergone a wide variety of neutral losses (**Figure 2c**). This indicates that considering appropriate neutral losses can again boost the quality of spectrum annotations.

### Code availability

spectrum_utils is available for Python 3.8+ and can be easily installed from the Python Package Index (PyPI) using pip or via conda using the Bioconda channel.^42^ spectrum_utils depends on NumPy^43^ and Numba^44^ for efficient numerical computation, Pyteomics^1,2^ for peptide fragment ion mass calculations and USI resolving, fastobo^45^ and lark-parser for parsing ProForma annotations, matplotlib^46^ for static plotting, and Altair^47^ and Pandas^48^ for interactive plotting.

All code and detailed documentation on how to use spectrum_utils is freely available as open source under the Apache 2.0 license at https://github.com/bittremieux/spectrum_utils. Code for the example spectrum plot in Figure 1b is available as open source at https://gist.github.com/bittremieux/47be3b3b038efb1c5b320a136b1e2d47 and code for the MassIVE-KB peak interpretation use case is available as open source at https://gist.github.com/bittremieux/cf58d176bba81ee0af1810f3728d150c. The results presented here were obtained using spectrum_utils v0.4.0.

## Conclusions

spectrum_utils combines a high-level Python API to easily perform common MS data tasks with only a single line of code, such as annotating MS/MS spectra with their peptide labels after database searching or visualizing spectrum–spectrum matches from spectral library searching using mirror plots. Additionally, power users can expand upon the spectrum_utils functions and infinitely customize their results by writing surrounding Python code. As an example, we have demonstrated these aspects by annotating fragment ions for over 2 million MS/MS spectra and providing insights into which ion types explain the observed peaks. This functionality is similar to the recently proposed MS_Piano software,^49^ but is inherently flexible using only a few lines of Python code, supports an extensive variety of PTMs through modification CV support in ProForma 2.0, and is fully cross-platform and open source. Further downstream applications can easily be achieved, such as quality assessment of spectrum annotations based on explained ion intensity or exporting annotated spectra to create a spectral library.

Besides the novel capabilities introduced here, the prior core functionality of spectrum_utils has also been extended and made compatible with third-party Python MS libraries focusing on both proteomics and metabolomics. Harnessing spectrum_utils’ plotting functionality to create publication-quality spectrum plots and mirror plots, this now powers on-the-fly spectrum visualization in the Metabolomics USI interface,^14^ for MS/MS spectra in the MassIVE web interface, and it is integrated into the Python software packages Pyteomics,^1,2^ pyOpenMS,^3^ and matchms^4^ for MS-based proteomics and metabolomics. In conclusion, spectrum_utils powers a growing open-source Python software stack and has emerged as a focal point of MS data manipulation for proteomics and metabolomics.

## Acknowledgements

This research was supported by BBSRC-NSF award 2152526 and National Institutes of Health award R03OD034493.

